# *CEL*eidoscope: quad-fluorescent *Caenorhabditis elegans* strain for tissue-specific spectral single-cell analyses

**DOI:** 10.64898/2026.03.25.714250

**Authors:** Clair R. Henthorn, Natalia Betancourt Rodriguez, Zach Stenerson, Kathy Vaccaro, Mostafa Zamanian

## Abstract

Cell and tissue-specific transcriptomic profiling of *Caenorhabditis elegans* is commonly achieved by fluorescence tagging or staining of targeted cell populations, often followed by fluorescence-activated cell sorting (FACS) and RNA sequencing. However, these approaches typically require separate strains for each labeled population, increasing labor and experimental variability while limiting direct comparison of multiple tissues within the same genetic background. To address this limitation and establish proof of concept, we engineered *CEL*eidoscope, a multicolored *C. elegans* strain that enables spectral sorting of multiple major cell types within a single strain population. Strain construction was carried out using a high-throughput screening method that reduces the labor and plastic costs associated with transgene integration and outcrossing. Four primary tissues (body muscle, neurons, intestinal, and pharyngeal muscle cells) were tagged with spectrally distinct fluorescent proteins, allowing compatibility with viability and nucleic acid dyes. Using spectral flow cytometry, dissociated *CEL*eidoscope cell suspensions could be sorted based on their spectral profiles, with cell recovery rates approximating the expected cell counts in whole organisms. Transcriptomic analysis of the sorted cell populations further validated the identity of the sorted populations, with recovered cells exhibiting gene expression signatures consistent with their intended cell and tissue identities. Together, these results establish *CEL*eidoscope as a versatile tool for multiplexed cell-type isolation in *C. elegans*, providing a framework for tissue-specific analyses from a common strain background.

## Introduction

The free-living nematode *Caenorhabditis elegans* has been used as a model organism in a wide range of biological research endeavors due to its amenable characteristics such as a transparent body, short life cycle, and simple anatomy (1,2). Its well-mapped genome and complete cell lineage have improved our understanding of metazoan developmental biology, neurobiology, and gene function and regulation (3–5). *C. elegans* has also served as a critical model of the study of some of the most prominent human diseases, including neurodegenerative diseases, diabetes, and cancer (6,7), as well as for studying conserved biological processes relevant to parasitic nematodes of plants and animals (8–12).

Despite its relatively simple anatomy, *C. elegans* exhibits extensive cellular specialization, with distinct tissues performing coordinated physiological functions. As a result, biological responses measured at the whole-organism level often reflect the combined action of multiple cell types, masking cell-specific transcription factors and regulatory mechanisms. Tissue-specific analyses are essential for understanding how gene expression and cell-signaling contribute to organism-level phenotypes.

Interest in analysing tissue-specific gene expression in *C. elegans* has increased in recent years, especially as it relates to the study of metazoan biology, the onset of diseases, and responses to chemical and environmental perturbations (13,14). There has been significant growth in methods and platforms to measure whole-organism phenotypes (15,16) and profile *C. elegans* and related parasitic nematodes at single-cell resolution (17–31). These methods either offer a broad summary of phenotypic outcomes or molecular insights into a small subset of cells, but they are unable to sufficiently connect the cellular and molecular roles of multiple tissue types to whole-organism phenotypes. A method that can bridge the gap for investigating biological responses at multiple levels of organization can help resolve questions relating to system-level biology.

To achieve this, we develop and validate *CEL*eidoscope, a multicolored *C. elegans* strain that enables multiplexed, tissue-resolved analysis from a single genetic background using spectral flow cytometric approaches. This strain expresses spectrally distinct fluorescent proteins driven by tissue-specific gene promoters located in primary somatic cell types, including neurons, body muscle, pharyngeal muscle, and intestinal cells. Early validation efforts have shown that *CEL*eidoscope enables unique identification and sorting of specific cell types using spectral flow cytometry, imaging, and cell sorting for RNA-sequencing. We further discuss the utility of this strain as a flexible method for integrating cell-type-specific and organism-level analyses across a wide range of biological applications.

## Results

### Construction of *Caenorhabditis elegans* strains with spectrally distinct cell types

We designed a *C. elegans* strain that would accommodate multimodal, simultaneous analysis of major somatic tissues using tissue-type specific promoters to drive the expression of spectrally distinct fluorescent proteins. The *C. elegans* hermaphrodite consists of five major tissues: epithelial, alimentary, muscle, neuronal, and reproductive. Its somatic cell lineages include 959 nuclei, with 302 neurons, 125 body muscle cells, 20 pharyngeal muscle cells, and 20 intestinal cells (32–35). Tissue-specific promoters were required to meet the following criteria: 1) confined expression to the target cell type, 2) constitutive expression in terminally differentiated cells, and 3) strong expression levels.

Body muscle cells, including those of the body wall, enteric, anal, vulval, and additional reproductive muscles, collectively express *myo-3*, which encodes for a myosin heavy chain protein of thick filaments in muscle cells (28,35–37). A second myosin heavy chain gene, *myo-2*, is exclusively expressed in striated muscle cells within the pharynx (32,38). Intestinal cells constitutively express *vha-6*, a gene encoding for a proton-translocating α-subunit of a vacuolar-type ATPase localized in the apical membrane (39). Several *C. elegans* pan-neuronal genes have also been demonstrated to have expression in other non-neuronal cell types, thus we chose the conservative promoter for *rgef-1*, which is expressed in 300/302 total neurons excluding the two CAN neurons (40,41). These four tissue-specific promoters (*myo-3*p, *myo-2*p, *vha-6*p, *rgef-1*p) were chosen to drive the expression of spectrally distinct fluorescent proteins previously validated for expression in *C. elegans* (**Figure 1A**) (42,43). Fluorescent proteins in the UV and far red spectra were omitted to allow for the incorporation of dyes used for markers of viability and/or nucleic acids.

**Figure 1.**
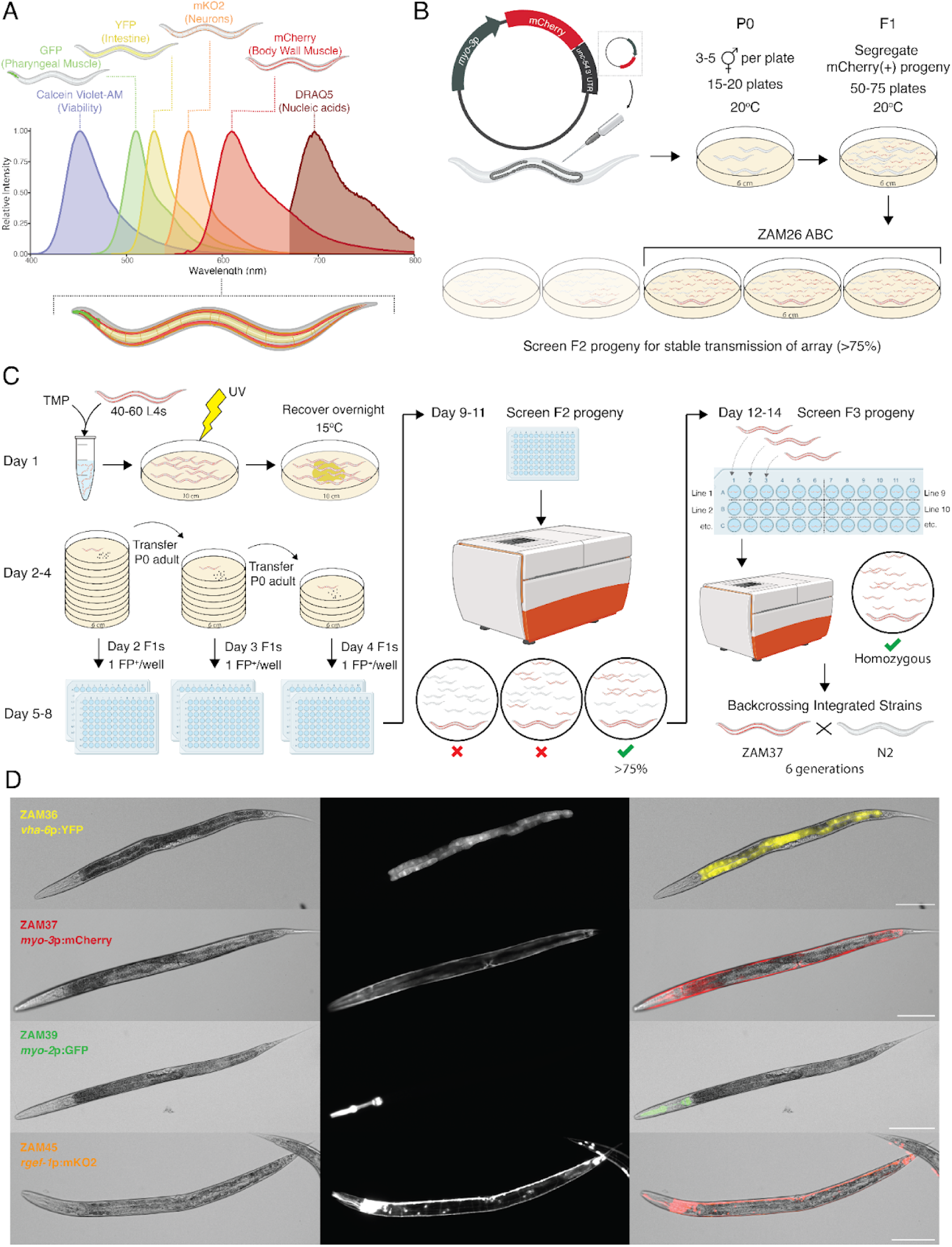
Spectral design and construction of *CEL*eidoscope constituent lines. (A) Emission profiles for four spectrally distinct fluorescent proteins used to tag specific cell types in *C. elegans* in combination with Calcein Violet-AM for cell viability and DRAQ5 to identify nucleic acids. (B) Extrachromosomal arrays with cell-type specific promoters driving expression of a fluorescent protein were microinjected into nonfluorescent *C. elegans* strain N2 worms. Progeny (F1 generation) were segregated to NGM plates and the F2 generation was evaluated for specific and complete fluorescent protein expression in the targeted tissue/cell-type. Lines were established in triplicate using progeny populations that exhibited strong and stable transmission (>75% progeny with positive fluorescence). (C) Extrachromosomal array integration to establish a 100% transmission rate of the transgene was facilitated by TMP-UV integration. 50-70 L4 stage transgenic worms were incubated in M9 medium containing trimethylpsoralen and then exposed to UV irradiation prior to incubation overnight at 15°C with *Escherichia coli* OP50 (Day 1). Next day, surviving worms were segregated to individual 6cm NGM plates for brood laying. Any surviving adult worms were transferred to a fresh 6cm NGM plate every day for 3 days to segregate progeny laid on days one, two, and three (Day 2-5). Fluorescent F1 progeny were picked into 96-well plates (∼150-200 L4s) containing growth media and incubated at 20°C with shaking until the F2 brood was laid (Day 6-9). All wells of each 96-well plate were screened for fluorescent progeny using the high-content imaging system. Wells where >75% of F2 progeny were fluorescent positive were identified as potential integration events (Day 10-12). Six F2 progeny from each independent integration event were segregated into individual wells of a 96-well plate containing growth media to screen for homozygous fluorescent F3 populations (Day13-15). (D) *C. elegans* strains expressing tissue-specific fluorescent proteins in the intestine (YFP), body muscle (mCherry), pharyngeal muscle (GFP), and neurons (mKO2; pan-neuronal). Scalebar=100µm.

Independent transgenic *C. elegans* strains were generated by microinjecting extrachromosomal arrays carrying the promoter::fluorescent protein constructs (**Figure 1B**). The F1 offspring from microinjected parents were screened for broods where at least 75% of the F2 generation expressed the transgene of interest and exhibited bright and complete cell-type expression. Each of the four independent fluorescent strains were established in triplicate to maintain genetic diversity. To ensure dependable transmission of the cell-type specific labels in each strain, extrachromosomal array integration into the *C. elegans* genome was completed using chemical mutagenesis and ultraviolet light radiation (**Figure 1C**).

Extrachromosomal array integration in *C. elegans* has, historically, been a reagent and labor intensive effort with significant challenges. Random integration of the array using paired chemical and UV radiation may result in unintended growth defects, lethality, and transgene silencing, and there is difficulty in distinguishing successful integration events from high transmission of residual extrachromosomal arrays. Additionally, screening mutagenized populations with low frequency integration events requires several hundreds *C. elegans* growth media plates, leading to excessive plastic use and labor-intensive maintenance (44–46). To circumvent these challenges, we developed a 96-well plate-based method for screening TMP/UV mutagenized progeny using liquid culture and fluorescence-based microscopy (**Figure 1C**).

Each transgenic fluorescent strain was subjected to the chemical mutagen TMP followed by exposure to UV radiation, as per established methods (45), and surviving parental worms were segregated to individual nematode growth media (NGM) plates (approx. 60-75 hermaphrodites). Over the course of three days, surviving hermaphrodites were transferred to fresh NGM plates (6 cm) to capture F1 progeny one, two, and three days after mutagenesis. We singled F1 progeny exhibiting bright, cell-type specific fluorescence (150-200 total worms per day) to individual wells of a 96-well plate containing liquid growth media supplemented with *Escherichia coli* OP50 (**Figure 1C**). This method bypassed the use of over a hundred NGM plates and allowed for the maintenance and tracking of more independent lines than the traditional method. The 96-well plates were screened with a high-content imager after the F2 brood was produced, and wells where the majority of F2 progeny (≥75%) were fluorescent were considered prospective integration events and propagated to identify homozygous lines. Similarly, six F2 progeny from each identified well were singled to individual wells of a 96-well plate for liquid culture to screen F3 populations. Wells where 100% of the progeny expressed the desired tissue-type-specific fluorophore were considered homozygous lines (**Supplemental Figure 1**) and selected for continued maintenance and observation on solid NGM plates. Any strains exhibiting slow growth, atypical morphology, or had integration events on the same chromosome as an already established strain were omitted and remaining homozygous lines were outcrossed with *C. elegans* N2 for six total generations to reduce undesired mutations induced by the integration process. Using this liquid growth media screening method, we generated *C. elegans* strains with the following genetically encoded fluorescent profiles: YFP^+^ intestinal cells (ZAM36), mCherry^+^ body muscle cells (ZAM37), GFP^+^ pharyngeal muscle cells (ZAM39), and mKO2^+^ pan-neuron cells (ZAM45) (**Figure 1D**).

### Outcrossing fluorescent strains to generate the quad-fluorescent *CEL*eidoscope strain

To enable multicellular, tissue-specific analyses and single-cell applications focusing on distinct cell populations, we combined our individual fluorescently tagged *C. elegans* strains into a single, quad-fluorescent worm strain. This multicolored strain was constructed by subsequently mating the single-color fluorescent worm strains (**Figure 2A**). This required every promoter-driven fluorescent protein in each strain to have randomly integrated on a different chromosome. Our approach utilized a traditional mating scheme beginning with ZAM36 (*vha-6*p::YFP) hermaphrodites and ZAM39 (*myo-2*p::GFP) males to produce dual fluorescent F1 progeny. All F1 hermaphrodite progeny expressing both GFP and YFP were mixed and the subsequent dual fluorescent F2 progeny were segregated into 96-well plates for liquid culture as previously described. The F3 generation was screened for independent populations where all progeny within a well were expressing both YFP and GFP fluorophores, indicating a homozygous line for both fluorophores. Hermaphrodites from the established dual fluorescent line (ZAM44) were subsequently mated with males with mCherry-tagged body muscle cells (ZAM37) and screened in a like manner to produce the tri-fluorescent ZAM46. The final neuronal-specific fluorophore, *rgef-1*p::mKO2::*unc-54* 3’ UTR, was incorporated in a similar manner to produce the final strain, *CEL*eidoscope (**Figure 2B**).

**Figure 2.**
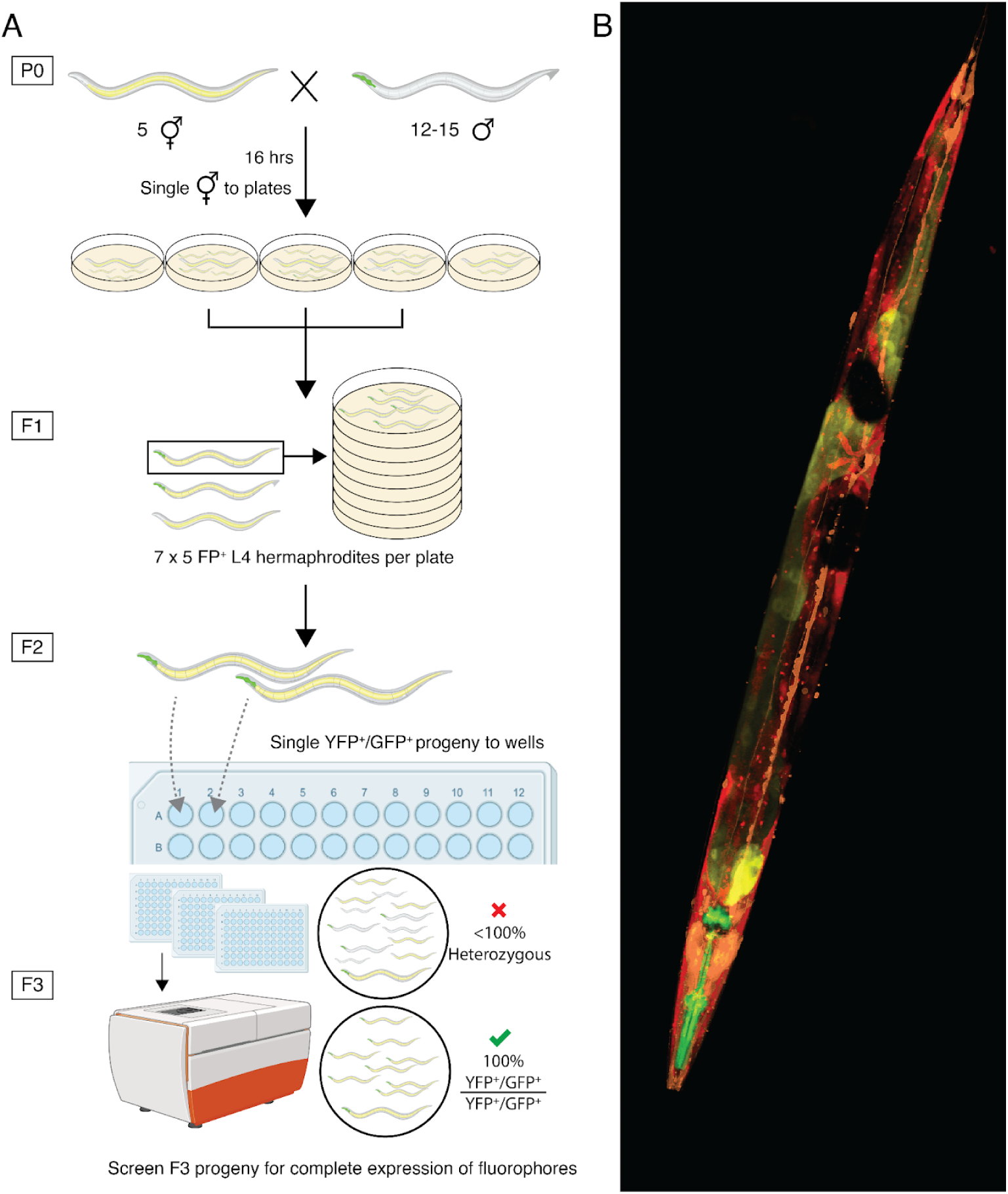
*\* assembly facilitated by *C. elegans* strain mating. (A) Schematic of mating scheme to assemble the *CEL*eidoscope worm strain. Combining the fluorophores through *C. elegans* mating required every promoter-driven fluorophore to have randomly integrated on a different chromosome. The integrated array producing ZAM36(*vha-6*p::YFP) is located on the X chromosome. Therefore all of the fluorescent markers were consolidated into a single *C. elegans* strain by subsequent mating of fluorescent males to YFP expressing hermaphrodites. Five YFP-hermaphrodites were mated with 12-15 fluorescent males (GPF/mCherry/mKO2) and co-incubated for 16 hrs and then hermaphrodites were segregated to individual 6 cm NGM plates to identify hermaphrodites that had successfully mated. Successful mates produced F1 progeny with the combined fluorescent profiles (YFP/GFP), which were propagated to new NGM plates (5 L4s/plate for 7 total plates). The F2 population was segregated to individual wells of a 96-well plate (200-384 total L4s) containing growth media and the F3 progeny were screened for wells where all progeny were expressing the desired fluorophores. Homozygous lines were maintained and mated with fluorescent males containing the next fluorophore to be added. (B) Confocal microscopy of young adult *CEL*eidoscope worm expressing all four tissue-specific fluorescent proteins.

The fluorophores used to construct *CEL*eidoscope cannot be distinguished by traditional microscopy methods due to the overlapping emission spectra among GFP, YFP, mCherry, and mKO2. Therefore, we exploited the spatial expression patterns of the fluorescent proteins to identify the presence or absence of a homozygous population using a high-content imaging system (**Supplemental Figure 2**). The final strain, *CEL*eidoscope (ZAM47), incorporates all four of the fluorescent translational reporters in a single strain.

### Validating the *CEL*eidoscope strain by spectral flow cytometry

We conducted a spectral flow cytometry pilot run to assess the functionality of *CEL*eidoscope for quantitative gene expression differences in cell populations. In order to successfully identify the cell-specific fluorescent populations in dissociated *CEL*eidoscope cell suspensions, we had to first validate our single-color *C. elegans* strains as technical controls for instrument set up and to generate the spectral unmixing matrix (**Figure 3A**). Negative technical control cell suspensions to account for cellular autofluorescence were generated using the non-fluorescently tagged *C. elegans* N2 strain. A portion of the un-tagged N2 cell suspension was stained with either DRAQ5 or Calcein Violet-AM to produce positive, single stain technical controls identifying nucleated and viable cells, respectively. Initially, single-stain controls for each fluorophore were produced by dissociating developmentally synchronized L4 populations from individual fluorescent strains (ZAM36, ZAM37, ZAM39, and ZAM45). However, due to the low abundance of certain fluorescent populations, particularly GFP⁺ and YFP⁺ cells, we incorporated commercially available single-color compensation beads for mCherry, YFP, and GFP to define the spectral matrix. These beads replaced the need to generate dissociated cell suspensions for the corresponding strains, substantially reducing preparation time. Because no compensation beads were available for mKO2, dissociated cells from the ZAM45 strain were retained for this channel. The final spectral matrix was therefore generated using a combination of compensation beads and single-color cell suspensions. After spectral unmixing, we analyzed multicolor panel controls that included the nucleic acid and viability dyes in each cell-type specific fluorescent cell suspension to optimize the gating strategy of each fluorophore prior to assessing the *CEL*eidoscope cell suspension (**Supplemental Figure 3**).

**Figure 3.**
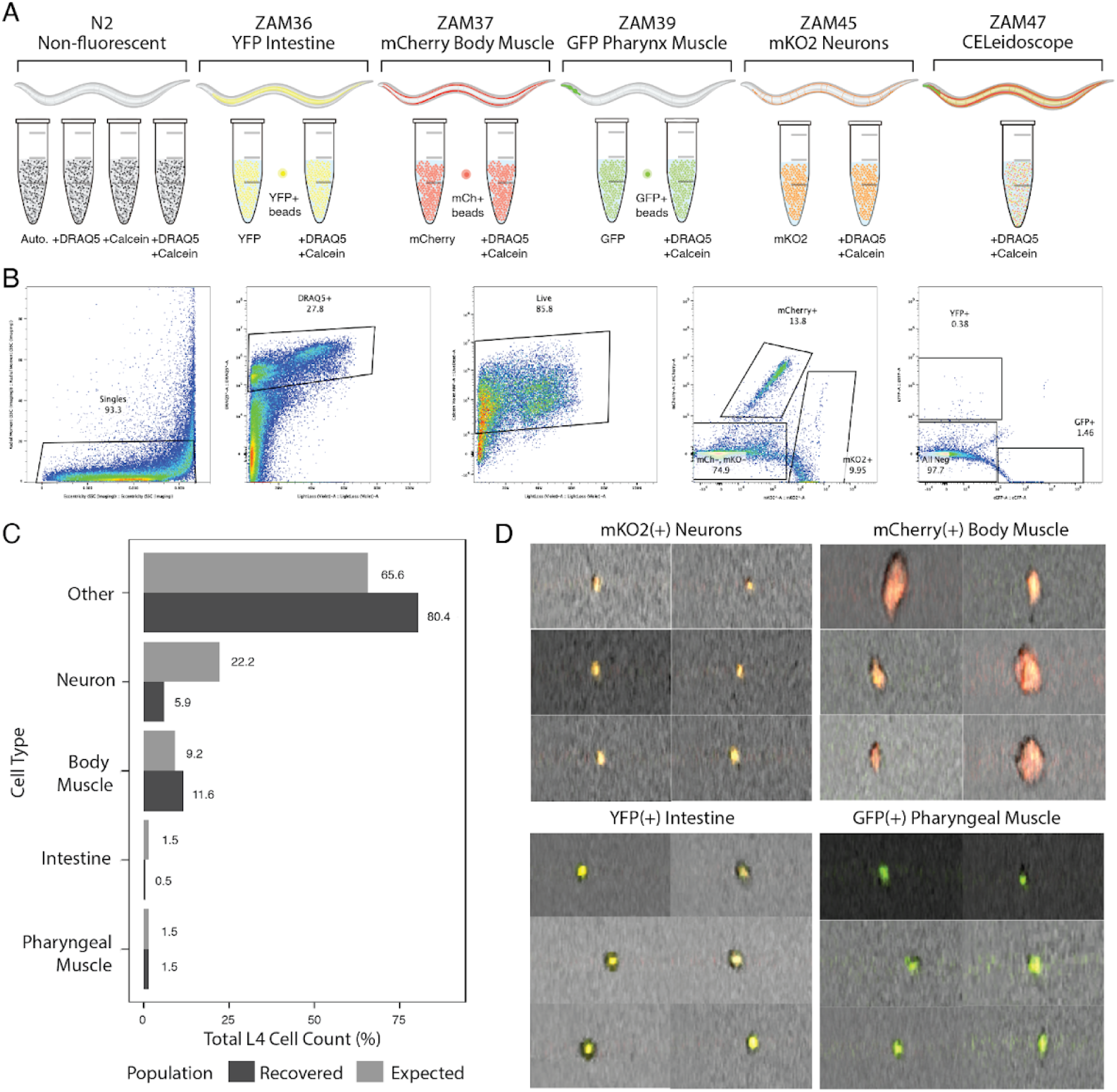
Spectral flow cytometry characterization of *CEL*eidoscope. (A) Schematic showing the worm strains and cell suspensions used to define the spectral matrix prior to analyzing the *CEL*eidoscope cell suspension. Non-fluorescent N2 cell suspensions were used as autofluorescence controls as well as single-color technical controls for viability (Calcein Violet-AM) and nucleic acid-containing objects (DRAQ5). The individual fluorescent strains were used to define the YFP, mCherry, GFP, and mKO2 spectral profiles. Additional cell suspensions containing viability and nucleic acid dyes were sorted to compare gene expression profiles with sorted cell suspensions obtained from *CEL*eidoscope suspensions. (B) Gating strategy based on single-color populations used to identify and sort *CEL*eidoscope cell populations. (C) Comparison of expected and recovered cell populations based on known cell abundances in L4 stage *C. elegans*. (D) Imaging of the cell populations show small round neurons, large and spindle-shaped body muscle cells, large round intestine cells, and small oblong pharynx muscle cells.

We next analyzed the *CEL*eidoscope cell suspension, applying the gating scheme set by the individual fluorescent multicolor technical control suspensions and compensation beads (**Figure 3B**). We sorted single cell populations that were nucleated and viable (DRAQ5^+^, Calcein Violet-AM^+^) and differentiated by the expression of mCherry, mKO2, YFP, GFP, or no fluorescence. We quantified the success of cell-type identification based on the L4 stage larvae containing 959 somatic cells, 400 germ cells, and assuming the dissociated cell suspensions reflected the biological composition of whole-tissue *C. elegans.* Neuronal and body muscle cells account for 22.2% and 9.2% of the total L4 cell count, respectively, while intestine and pharyngeal muscle account for 1.5% each (2,47). Cell sorting of the gated populations recovered 5,796 neuronal cells and 11,404 muscle cells, representing approximately 5.9% and 11.6%, respectively, of the total cell population (98,171 total cells recovered) (**Figure 3C**). A total of 502 intestinal cells and 1,498 pharyngeal muscle cells were recovered, representing 0.5% and 1.5% of the total cell population, respectively (**Figure 3C**). Imaging analysis of the mKO2^+^ population shows orange, round, and small cells, representative of neuron morphology in cell suspensions (48). The mCherry^+^ population shows red, large, more oblong cellular objects, reflective of the spindle-shape morphology of *C. elegans* body muscle cells (**Figure 3D**). The YFP^+^ population contains yellow, round cells resembling intestinal cells, and the GFP^+^ population exhibits green, small, more oblong cells typical of pharyngeal muscle cells (**Figure 3D**).

### Transcriptomic validation of FACS-defined *CEL*eidoscope populations

To independently validate the enrichment and recovery of the intended tissues through FACS, we performed bulk RNA sequencing on each of the sorted cell populations. This analysis aimed to assess three features: 1) expression of endogenous genes associated with each promoter, 2) alignment of sequencing reads to the corresponding fluorescent protein, and 3) enrichment of broader tissue-specific markers. Promoter-associated genes showed enrichment in their expected populations, particularly in the mKO2^+^, mCherry^+^, and GFP^+^ populations (**Figure 4A**). Expression of the intestinal marker *vha-6* was minimal across populations, with low-level expression in the mKO2^+^ population. Notably, the YFP^+^ population exhibited low expression of the selected tissue-specific genes, but showed high expression of other intestinal markers such as *act-5*, *ifb-2*, and *mlt-2*, suggesting either incomplete recovery of promoter-driven transcripts or potential silencing associated with the integrated array.

**Figure 4.**
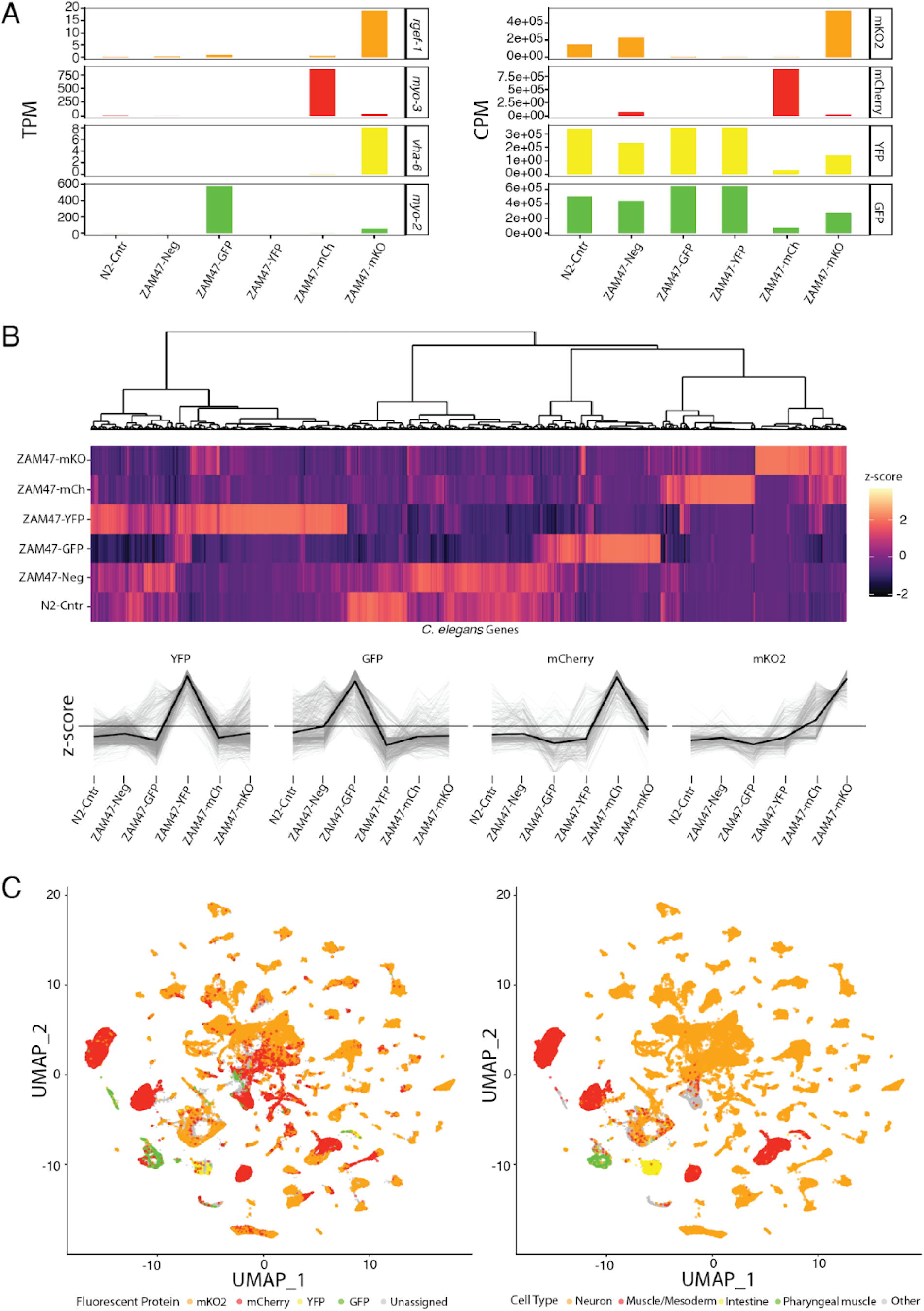
Transcriptomic analysis of *CEL*eidoscope fluorescent populations. (A) Expression of tissue-specific genes (TPM; left) and read alignment to fluorescent protein sequences (CPM; right) across *CEL*eidoscope samples. Tissue markers validate population enrichment, and fluorescent protein alignment confirms construct expression. (B) Hierarchical clustering based on z-score-normalized expression of most expressed *C. elegans* genes. Clusters 4, 5, 1, and 6 show preferential upregulation in YFP^+^, GFP^+^, mCherry^+^, and mKO2^+^ populations, respectively. (C) Projection of population-enriched gene clusters onto the CeNGEN single-cell *C. elegans* reference atlas UMAP using module score analysis (left). Corresponding annotated cell types are shown (right), demonstrating enrichment in neuronal, muscle/mesodermal, intestinal, and pharyngeal muscle populations.

To assess fluorescent protein specificity, sequenced reads were aligned directly to mKO2, mCherry, YFP, and GFP transcripts and normalized to counts per million (CPM). Each fluorescent population demonstrated the highest CPM values for its corresponding fluorescent protein, with strongest specificity observed in the mKO2^+^ and mCherry^+^ populations (**Figure 4A**). Some cross-alignment between YFP and GFP was observed, consistent with their high sequence similarity (>94% sequence identity) and spectral overlap.

To assess gene expression patterns, genes were hierarchically clustered based on z-score-normalized expression (**Figure 4B**). Partitioning the dendrogram into six clusters identified distinct gene sets enriched within each fluorescent population. Four clusters exhibited strong population-specific expression and were projected onto the CeNGEN single-cell UMAP (25) to assess correspondence with annotated cell types (**Figure 4C**). Genes enriched in the mKO2^+^, mCherry^+^, YFP^+^, and GFP^+^ populations localized predominantly to neuronal, muscle/mesoderm, intestinal, and pharyngeal muscle clusters, respectively. While some overlap was observed between spectrally similar fluorophores (mCherry and mKO2), overall consensus between fluorescence-defined populations and CeNGEN-annotated tissues supports enrichment of the intended cell types. Increasing clustering resolution beyond six clusters resulted in additional subdivision of populations without revealing new tissue-specific identities, suggesting that the six-cluster model most accurately captures meaningful distinctions among sorted populations.

## Discussion

Since the development of single-cell dissociation protocols in embryonic, larval, and adult stage worms, several cell-type-specific transcriptome libraries have been produced, including comprehensive transcriptomic atlases across various developmental stages, leading to novel cell-type-specific markers, regulatory elements, and insights into developmental and systems biology (20–22,24,25,27). Even though high-throughput single-cell methodologies exist and are highly informative, cell-type-specific methods that utilize economic technologies can be more flexible and accommodate diverse technical assay endpoints. Flow cytometry-based techniques that rely on a fluorescently-tagged cell population, usually using single-color GFP tagged strains, continue to be used to answer tissue specific hypotheses or enrich for particular cell populations intended for downstream cell culture assays (18,23,25,30,48–54). However, these approaches have focused on single-tagged strains, isolating one cell or tissue type at a time, and requiring multiple independent strains to examine different cell types in *C. elegans. CEL*eidoscope improves upon this by using four distinct fluorescent proteins to identify major tissue types within *C. elegans*, enabling simultaneous analysis of these cell types in response to various stimuli. By combining this quad-fluorescent strain with advanced single-cell technologies, such as spectral flow cytometry and imaging, *CEL*eidoscope allows for multimodal characterization, including viability, morphology, transcriptomics, and drug bioavailability, all from a common organismal population. To demonstrate its potential, we launched a pilot study to validate spectral flow cytometry as a method for profiling dissociated *CEL*eidoscope cell suspensions.

We began the construction of this new resource by creating independent *C. elegans* strains each expressing a unique fluorophore driven by a tissue-specific gene promoter in the form of an extrachromosomal array. Microinjection of the arrays into the *C. elegans* gonads is an effective way of producing transgenic progeny but the transmission of the array is transient and requires integration into the *C. elegans* genome for stable transmission. A well-known challenge in *C. elegans* gene manipulation methods (e.g., extrachromosomal array integration, CRISPR-Cas9, strain mating) is the labor-intensive and costly screening required to isolate a homozygous mutant of interest. This process often involves tracking independent progeny lines, requiring hundreds of NGM plates. Mariol et al. addressed this issue by reducing the number of NGM plates needed for extrachromosomal array integration from 2,094 to 498 by starting with parental lines that strongly transmitted the transgene of interest (46). However, high transmission lines can make it difficult to distinguish true integration events from high transmission rates, and this approach relies on a visible phenotype that may not be applicable to all methods of genetic manipulation. Our improved 96-well plate method facilitates the screening of more progeny with observable phenotypes and significantly reduces reliance on single-use plastics while maintaining independent progeny lines. Fluorescent progeny populations can be analyzed using either a plate-based imaging system or a simple microdissection scope with an LED excitation light system for bright, multicellular phenotypes. This method greatly reduces the time and cost associated with extrachromosomal integration and successfully identified a *CEL*eidoscope line homozygous for all four fluorescent reporters without requiring copious amounts of NGM plates.

We applied single-cell spectral flow cytometry paired with imaging and fluorescence-activated cell sorting to validate *CEL*eidoscope as a tool for simultaneous tissue-type analyses based on the cell-type-specific fluorescent profile. Assuming the L4 stage *CEL*eidoscope worms we dissociated into single-cell dissociations have 959 somatic and 400 germ cells (2,47), neurons should account for 22.2% of the cell population in the *CEL*eidoscope cell suspension. We recovered significantly fewer neuronal cells than expected (**Figure 3C**), which may be an outcome reflective of the population’s small cell size being similar to residual debris remaining in the cell suspension and therefore omitted by the applied gating strategy. In addition, dissociation conditions may contribute to neuronal underrepresentation: overdigestion may reduce viability and result in exclusion during live-cell gating, whereas underdigestion may lead to incomplete dissociation and exclusion during doublet discrimination. Low representation of FAC-sorted fluorescently-tagged neuronal cells has been observed in the past and rectified by using alternative gating approaches to isolate the target population (30). Despite the reduced overall recovery, we detected a broad diversity of neuronal cell types, including rare populations such as the DVA tail neuron. A refined gating approach guided by both expression of the mKO2 fluorophore and imaging capabilities of the spectral flow cytometer could improve the recovery rate of neuron populations.

In addition to neuronal cells, we also recovered fewer intestinal cells than expected from dissociated L4 stage *C. elegans*. Several factors likely contributed to the underrepresentation of this population. First, this population accounts for less than 3% of total cells in L4 stage *C. elegans,* potentially providing insufficient representation to define the GFP and YFP spectral profiles needed for spectral unmixing and *CEL*eidoscope analysis. Second, *C. elegans* exhibits intrinsic fluorescence in the GFP and YFP channels, specifically in the gut and gonads (55,56). Although we define gating parameters with YFP^+^ compensation beads and control for autofluorescence using non-fluorescent N2 cells, the low abundance of GFP- and YFP-labeled cells may not spectrally unmix correctly. Finally, intestinal epithelial cells are large and structurally polarized, making them more susceptible to damage or loss during enzymatic dissociation and filtration, which may further reduce their recovery in sorted populations.

*CEL*eidoscope is a powerful and versatile system that integrates high-resolution cellular characterization and whole-organism phenotyping. This dual approach enables the investigation of tissue-specific gene expression, developmental processes, and intercellular signaling within a unified experimental framework. By allowing multiple major tissues to be analyzed simultaneously from a single population, *CEL*eidoscope facilitates studies of cellular heterogeneity, coordinated tissue responses, and regulation under diverse biological conditions. Dissociated cells can be assessed for viability, morphology, and gene expression, and specific populations can be isolated for downstream analyses, including transcriptomics and functional assays. In addition, the platform is compatible with applications involving chemical or environmental perturbations, where tissue-specific responses can be resolved within a whole-organism context. *CEL*eidoscope’s utility extends beyond flow cytometry, as specific cell populations can be enriched through FACS and cultured in peanut lectin-coated chambers for physiological and/or biochemical assays, such as calcium flux measurements or ion flow monitoring in specific cell types. Overall, *CEL*eidoscope is a versatile tool with broad applications in both whole-organism and single-cell assays, offering the potential to enlighten our understanding of basic biology and tissue-specific transcriptomics that are otherwise inaccessible or challenging to study.

## Materials and methods

### *C. elegans* maintenance

*C. elegans* strains were maintained at 20°C on nematode growth media (NGM) agar plates seeded with *Escherichia coli* strain OP50. Strains were propagated by routine transferring of L4 stage worms to seeded NGM plates as previously described (57). Transgenic fluorescent larvae were identified using a NIGHTSEA^TM^ fluorescence adapter for a stereo microscope system (Electron Microscopy Sciences) equipped with the royal blue, cyan, or green filter set.

### Cloning expression plasmids

*C. elegans* tissue-specific promoters for the pharyngeal muscle, intestine, body muscle, and pan-neuronal cells were selected based on published literature and the *C. elegans* CeNGEN single-cell transcriptomic atlas (25). Promoter sequences were derived from WormBase Parasite (WBPS17) (58) targeting the 1.6 kb region upstream of the transcriptional start site. Fluorescent proteins with distinct spectral profiles were identified from prior *C. elegans* studies (42,43). All primers for promoter and fluorescent protein amplification are listed in **Table 1**. Promoter and fluorescent protein sequences were amplified using Q5 Taq polymerase (New England Biolabs, NEB) and purified using the QIAquick PCR purification kit (Qiagen). Promoter PCR products were digested with *HindIII* and *EcoRV* and fluorescent protein sequences were digested with *EcoRV* and *EcoRI* per the manufacturer’s instructions (NEB). The pPD95.75 plasmid backbone was prepared with double digestion using *EcoRI* and *HindIII*. Ligation of the backbone, promoter, and fluorescent protein sequences was performed with T4 DNA Ligase (NEB) per the manufacturer’s instructions. Ligated constructs were transformed into chemically competent *E. coli* DH5α cells (50 µL) by adding 2.5 µL or 5 µL of the ligation mixture to the competent cells and incubating on ice for 30 min prior to heat shock in a 42°C water bath for 45 sec followed by immediate induction on ice for 2 min. After incubation, 900 µL of SOC medium was added to the transformed cells and incubated at 37°C for 1 hr with shaking (200 rpm). The transformed cells were plated on LB agar containing ampicillin (100 µg/mL) and incubated at 37°C overnight. Cells containing the properly assembled construct were confirmed using colony PCR and plasmid sequencing. All arrays generated and used for this study are outlined in **Table 2**.

**Table 1.**
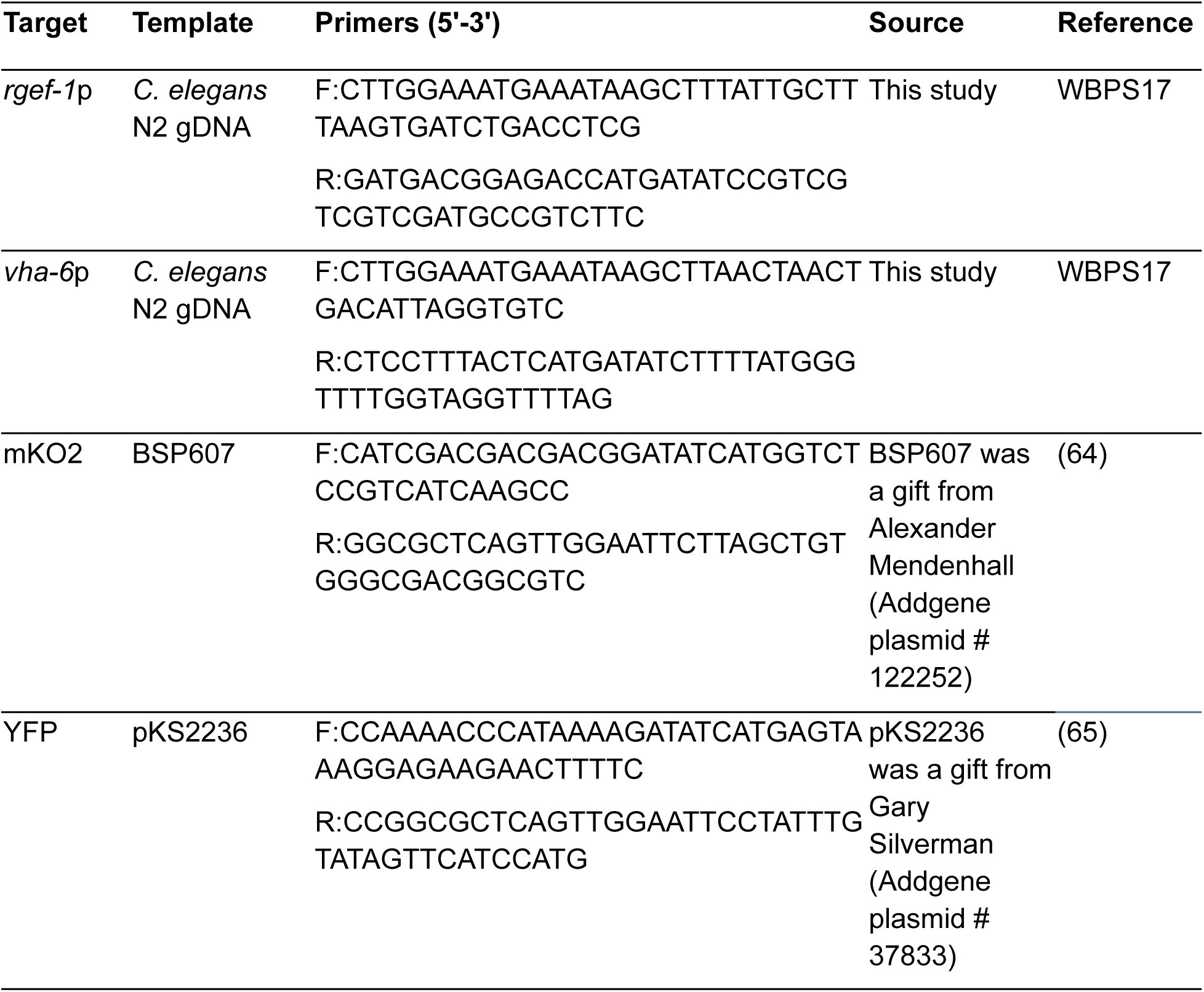
Primers used to generate the promoter and fluorescent protein constructs.

**Table 2.**
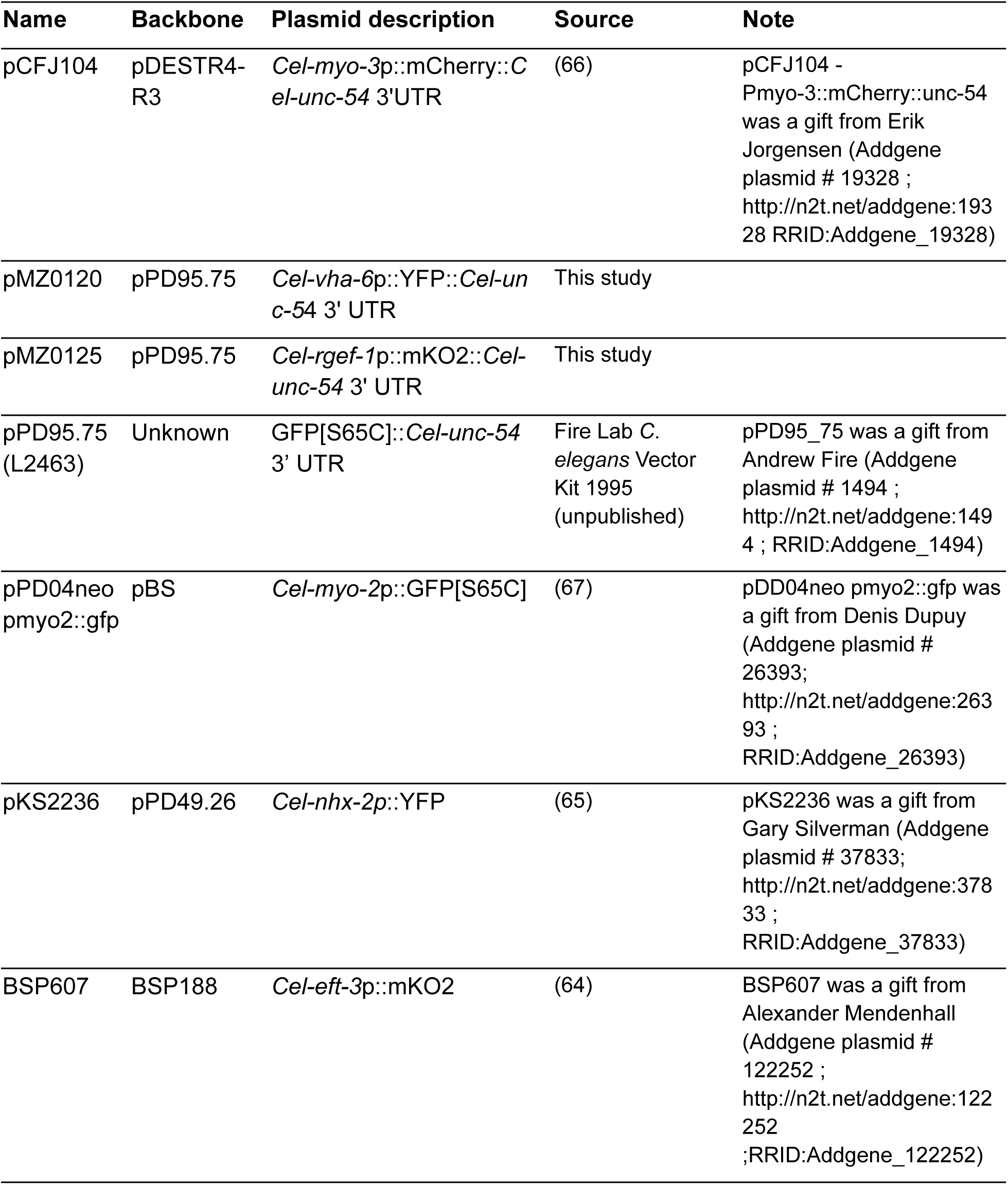
List of constructs generated or used in this study.

| Name | Backbone | Plasmid description | Source | Note |
| --- | --- | --- | --- | --- |
| pCFJ104 | pDESTR4-R3 | *Cel-myo-3p::mCherry::Cel-unc-54* 3'UTR | (66) | pCFJ104 - Pmyo-3::mCherry::unc-54 was a gift from Erik Jorgensen (Addgene plasmid # 19328 ; http://n2t.net/addgene:19328 RRID:Addgene_19328) |
| pMZ0120 | pPD95.75 | *Cel-vha-6p::YFP::Cel-unc-54* 3' UTR | This study |  |
| pMZ0125 | pPD95.75 | *Cel-rgef-1p::mKO2::Cel-unc-54* 3' UTR | This study |  |
| pPD95.75 (L2463) | Unknown | GFP[S65C]::*Cel-unc-54* 3' UTR | Fire Lab C. *elegans* Vector Kit 1995 (unpublished) | pPD95_75 was a gift from Andrew Fire (Addgene plasmid # 1494 ; http://n2t.net/addgene:1494 ; RRID:Addgene_1494) |
| pPD04neo pmyo2::gfp | pBS | *Cel-myo-2p::GFP[S65C]* | (67) | pDD04neo pmyo2::gfp was a gift from Denis Dupuy (Addgene plasmid # 26393; http://n2t.net/addgene:26393 ; RRID:Addgene_26393) |
| pKS2236 | pPD49.26 | *Cel-nhx-2p::YFP* | (65) | pKS2236 was a gift from Gary Silverman (Addgene plasmid # 37833; http://n2t.net/addgene:37833 ; RRID:Addgene_37833) |
| BSP607 | BSP188 | *Cel-eft-3p::mKO2* | (64) | BSP607 was a gift from Alexander Mendenhall (Addgene plasmid # 122252 ; http://n2t.net/addgene:122252 ;RRID:Addgene_122252) |

### *C. elegans* transgenesis

Transgenic *C. elegans* strains were created by microinjecting assembled arrays into the gonadal arms of *C. elegans* young adults. Injection mixes included 25 ng/µL of the transgenic plasmid, 75 ng/µL of pPD95.75 and nuclease-free water to a final volume of 30 µL. Microinjection were performed using a Zeiss AX10 microscope equipped with a 10x objective with glass 1BBL w.FIL 1.0 mm 4 IN filament (World Precision Instruments) injection needles assembled into a Narishige micromanipulator controlled by a FemtoJet (Eppendorf). Microinjected *C. elegans* were recovered on seeded NGM plates (6 cm) and maintained at 20°C. F1 progeny expressing tissue-specific fluorescence were monitored and lines with a high transmission rate (≥75%) were propagated at the L4 stage. Generated strains are listed in **Table 3**.

**Table 3.**
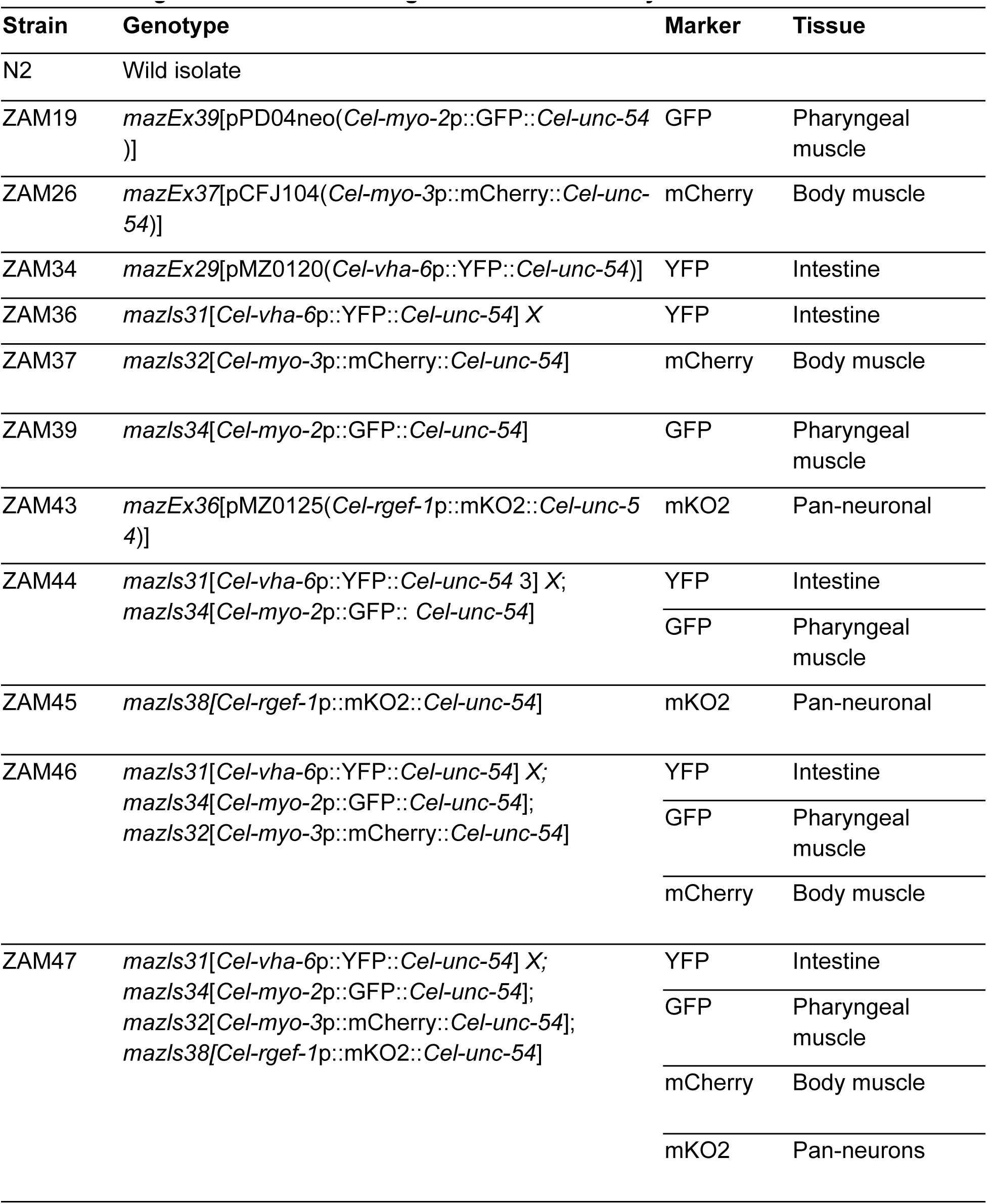
*C. elegans* strains used and generated in this study.

### Transgene integration

Stable chromosomal integration was achieved using trimethyl psoralen (TMP) and UV light radiation as previously described with modifications (45). *C. elegans* L4 stage worms expressing the transgene array (∼60-80 total worms) were incubated in M9 buffer (380 µL) containing 20 µL of TMP (1 mg/mL in dimethylformamide) for 15 min at room temperature protected from light on a rotating platform. After incubation, the worms in solution were transferred to an unseeded 10 cm NGM plate and the plate was exposed to long wave UV light radiation (350 µJ x 100) using the Stratalinker 1800. The plate was then seeded with 400 µL concentration *Escherichia coli* strain HB101 and incubated at 15°C overnight. Next day, any surviving worms were segregated to individual 6 cm seeded NGM plates and placed at 20°C. The surviving single, mutagenized worms were allowed to lay progeny over the course of three days, moving the parental worm to a new 6 cm seeded NGM plate every day. Once the progeny hatched and developed to L3/L4 stage, worms expressing the transgene were transferred to individual wells of a 96-well plate containing 200 µL K media supplemented with 50 µg/mL kanamycin and 2 µL of concentrated *E. coli* strain HB101. On average, 450-600 worms segregated across all three days of progeny. The plates were sealed with a breathable film and placed in a humidity chamber maintained at 20°C with shaking until the F2 generation hatched. Wells were ≥75% of progeny expressing the transgene of interest were transferred to 6 cm seeded NGM plates until larvae reached the L4 developmental stage. Six L4 stage worms from each independent line were segregated into individual wells of a 96-well plate as previously described to screen the F3 generation for a homozygous line (100% transgene expression). Wells displaying a homozygous fluorescent population were transferred to 6 cm NGM plates and maintained while confirming the stable integration of the transgene into the genome. Final homozygous lines were backcrossed to the parental background strain (*C. elegans* strain N2) six total generations to eliminate unintended background effects generated by the integration procedure.

### *C. elegans* strain mating

Strains expressing individual fluorescence markers were combined through the mating of the strains. For instance, five ZAM36 hermaphrodites were crossed with 12-15 ZAM39 males and allowed to copulate for 16 hrs at 20°C. Hermaphrodites were transferred to individual 6 cm NGM plates after 16 hrs to identify hermaphrodites that had successfully mated with ZAM39 males. F1 progeny expressing both GFP and YFP were transferred to 6 cm NGM plates (∼5 L4 transgenic worms per plate for 7 total plates) to lay the F2 generation. F2 progeny expressing both fluorophores were segregated into wells of a 96-well plate as previously described to screen for a homozygous line for both fluorophores in the F3 generation. Screening for the expression of both fluorophores was completed using the Molecular Devices ImageXpress Nano high content imaging instrument and applying the GFP and Cy3 filters. The sequential addition of tissue-specific fluorophores into one final strain was completed in the same manner using strains ZAM37 and ZAM45 to generate the complete quad fluorescent strain ZAM47.

### Single-cell dissociation

Developmentally synchronized L4 stage *C. elegans* were used for all single-cell dissociations. Animals for each strain were washed from the NGM plates into 15 mL conical tubes using M9 medium and washed until no *E. coli* OP50 remained in the suspension. The worm pellets were transferred to 1.5 mL Protein LoBind Tubes (Eppendorf) for a desired pellet size of ∼100 µL and dissociated as previously described (26). Undigested worm bodies were removed from the suspension by filtering through a 25 µm nylon mesh filter. The clean cell suspension was transferred to a new 1.5 mL protein LoBind tube and placed on ice for nucleic acid and viability staining. All cell suspensions were split so that each strain had a single-color cell suspension or no fluorescence for N2 autofluorescent technical control. For stained cell suspensions, Calcein Violet-AM (1 µg/mL) and/or DRAQ5 (1:2000 final dilution) was incubated with the cell suspensions for 20 min on ice in the dark. The cell suspensions were washed once by centrifuging at 1,000 x *g* for 6 min and resuspended in >300 µL M9 (340 mOsm/kg) + FBS (3%) for analysis.

### Spectral flow cytometry and sequencing

Flow cytometry was performed on a BD FACSDiscover^TM^ S8 spectral sorter, and data were analyzed with FlowJo^TM^ v10.10 Software (BD Life Sciences) (59) and the BD CellView Lens plugin for visualizing cell population imaging data. Single-cell suspensions were first gated to exclude debris and doublets based on forward (LightLoss (Violet)-A) and side scatter (SSC (Imaging)-A), followed by selection of nucleated and viable cells using DRAQ5 and Calcein Violet-AM staining, respectively. N2 control cells were used to establish the autofluorescence reference for spectral unmixing and served as the unstained control for gating. Spectral unmixing was established using single-color controls. Compensation beads labeled with mCherry, YFP, and GFP (Invitrogen) were used to define the corresponding spectral signatures, while the mKO2 signal was defined using the single-fluorophore strain ZAM45. Fluorescent cell populations were sorted using an 85 µm nozzle into TRIzol LS for RNA extraction using the Direct-zol^TM^ RNA MiniPrep kit (Zymo Research) following manufacturer’s specifications with optional DNase treatment step. Libraries were prepared using the low-input RNA workflow of the NEBNext Single Cell/Low Input RNA Library Prep Kit for Illumina (NEB) and sequenced on an Illumina NovaSeq X Plus to generate approximately 10 million paired-end reads (2 × 150 bp) per sample. Sequencing reads were quantified at the gene level and normalized to transcripts per million (TPM) using *C. elegans* gene lengths obtained from WormBase (release WBPS19). TPM values were filtered, z-score normalized across samples, and used for hierarchical clustering based on Euclidean distance with Ward’s method in R (v4.5.0) (60). Fluorescent reporter expression was quantified by aligning reads to protein coding sequences (mKO2, mCherry, YFP, GFP) using bwa (v0.7.19-r1273) (61) and samtools (v1.23) (62), and normalized to counts per million (CPM) based on total mapped reads per sample.

### Microscopy

Confocal microscopy of ZAM47 was performed on a Nikon A1R-Si+ confocal microscope equipped with six laser lines (405, 440, 488, 514, 561, and 640 nm) for multi-channel imaging, a 20X objective, and the Nikon Elements Analysis software. FIJI was used to collapse Z-stack projections for a maximum projection image of each wavelength (63).

## Data Availability

Sequencing data are available at NCBI SRA (PRJNA1442645). FlowJo data files are available in Zenodo (10.5281/zenodo.19221994). Scripts used in the processing and visualization of data are available at github.com/zamanianlab/celeidoscope-ms. The *CEL*eidoscope strain (ZAM47) will be deposited at the *Caenorhabditis* Genetics Center (CGC, University of Minnesota) and made publicly available upon publication.

## Acknowledgements

The authors thank the University of Wisconsin Carbone Cancer Center (UWCCC) Flow Cytometry Laboratory for use of its facilities and services. The authors also thank Christopher Miller for his help cloning constructs for extrachromosomal array expression in *C. elegans*.

## Supplemental Figures

**Supplemental Figure 1.**
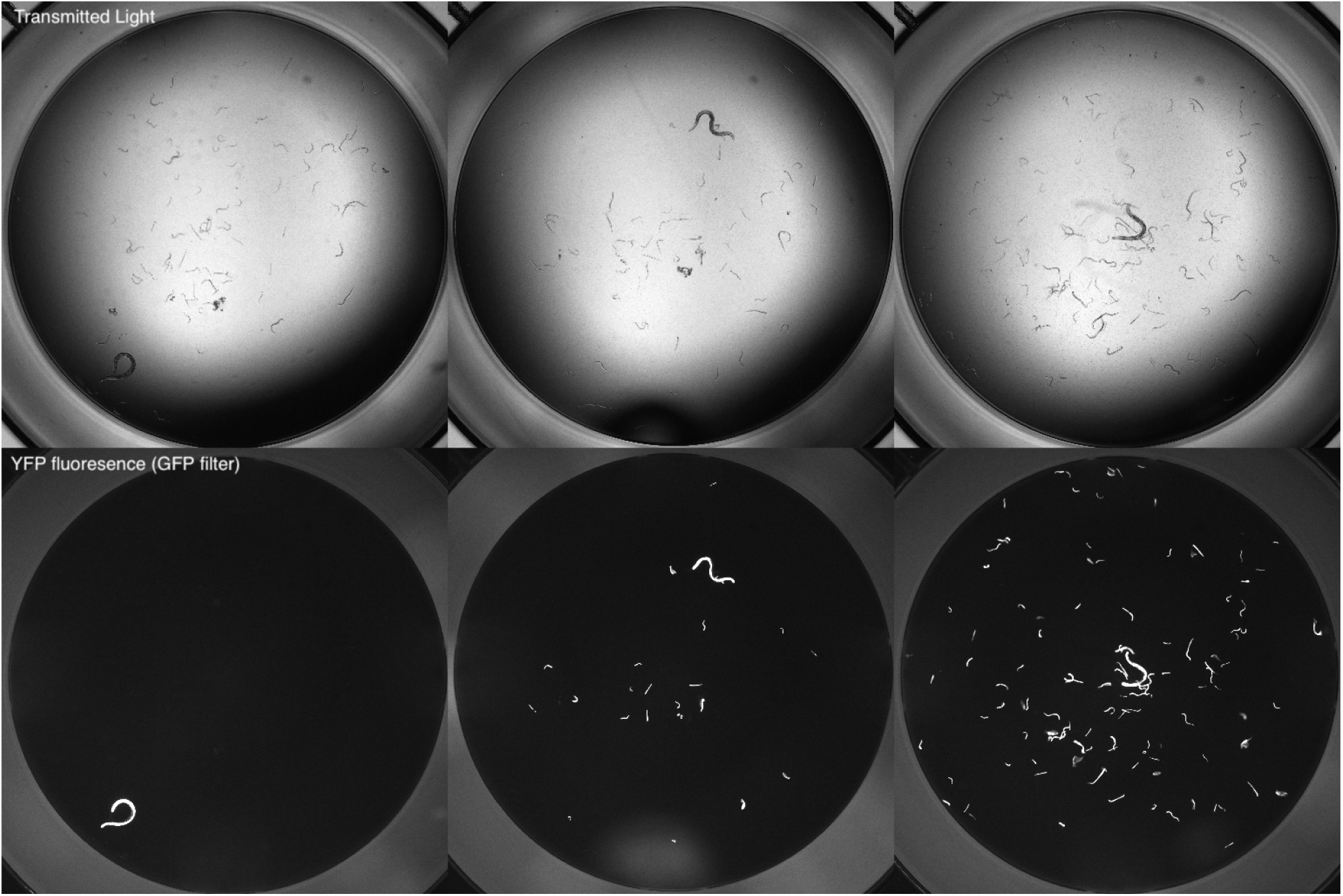
Images representing fluorescence screening of F3 progeny during the extrachromosomal array integration process. F2 larvae with YFP^+^ intestinal cells giving rise to no fluorescent progeny (left), partial YFP^+^ F3 population (middle), and complete YFP^+^ expression in the intestines of all F3 progeny (right).

**Supplemental Figure 2.**
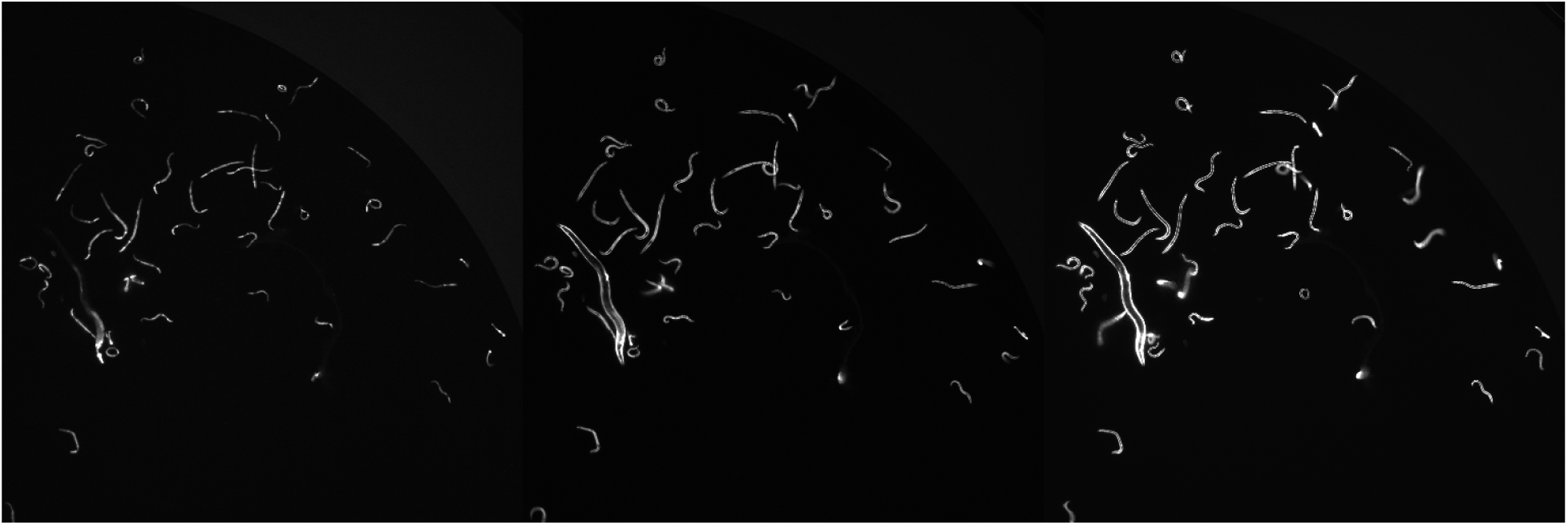
Representative images of multicolored *C. elegans* strain exhibiting fluorescence in GFP, CFP, and Texas Red filters of the ImageXpress Nano high-content imager. Single worm strain exhibiting GFP^+^ pharynx and YFP^+^ intestine in GFP channel (left), YFP^+^ intestine and mCherry^+^ body muscle cells in the CFP channel (middle), and mCherry^+^ body muscle cells in the Texas Red channel (right).

**Supplemental Figure 3.**
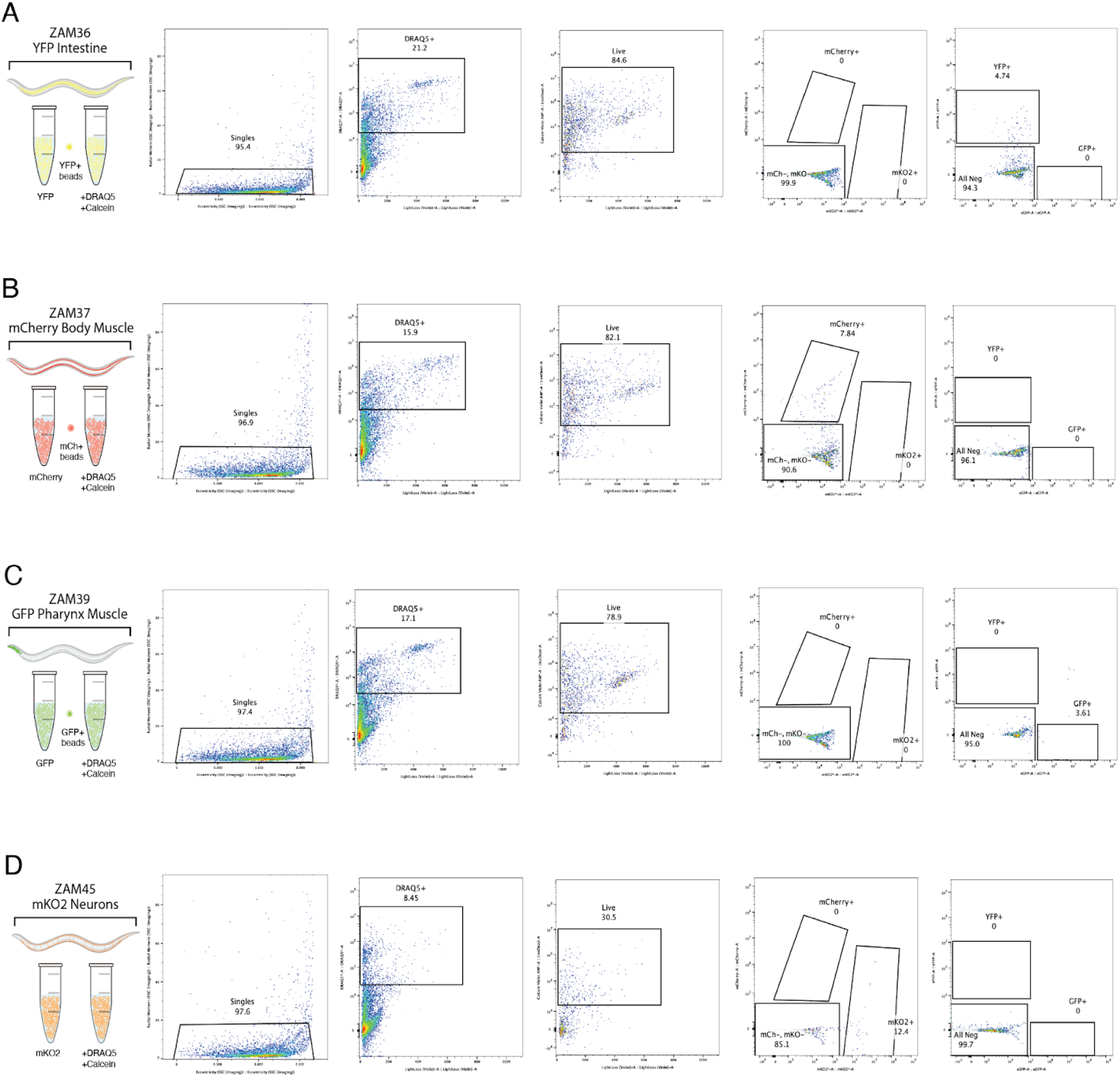
Validation of spectral unmixing matrix using single-fluorophore *C. elegans* strains. (A) Gating strategy defined for YFP^+^ cells using YFP^+^ compensation beads and ZAM36 strain. (B) Gating strategy defined for mCherry^+^ cells using mCherry^+^ compensation beads and ZAM37 strain. (C) Gating strategy defined for GFP^+^ cells using GFP^+^ compensation beads and ZAM39 strain. (D) Gating strategy defined for mKO2^+^ cells using ZAM45 strain.

