## Supplementary figures and images for "*CEL*eidoscope: quad-fluorescent *Caenorhabditis elegans* strain for tissue-specific spectral single-cell analyses"

### Supplemental Figure 1

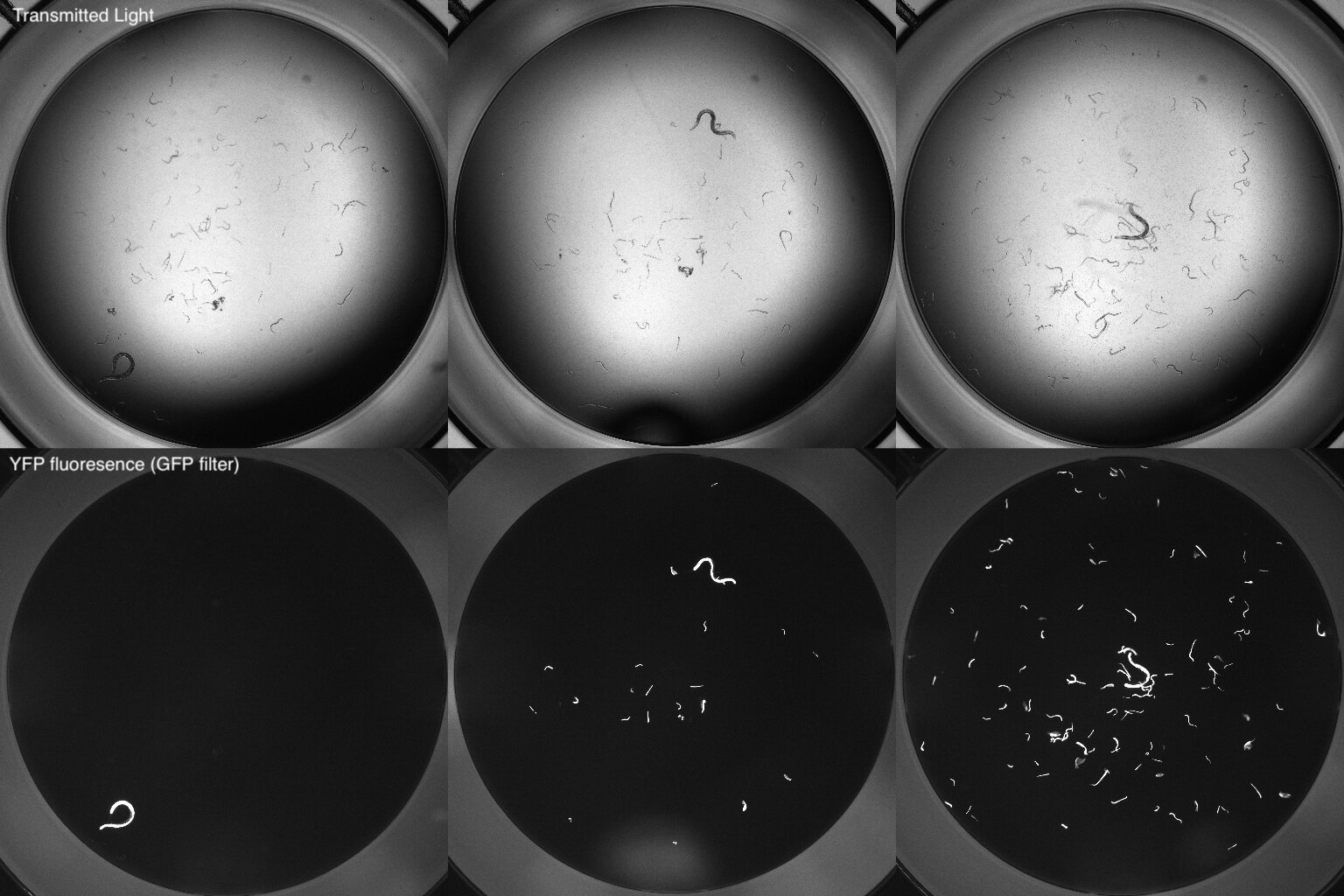

### Supplemental Figure 2

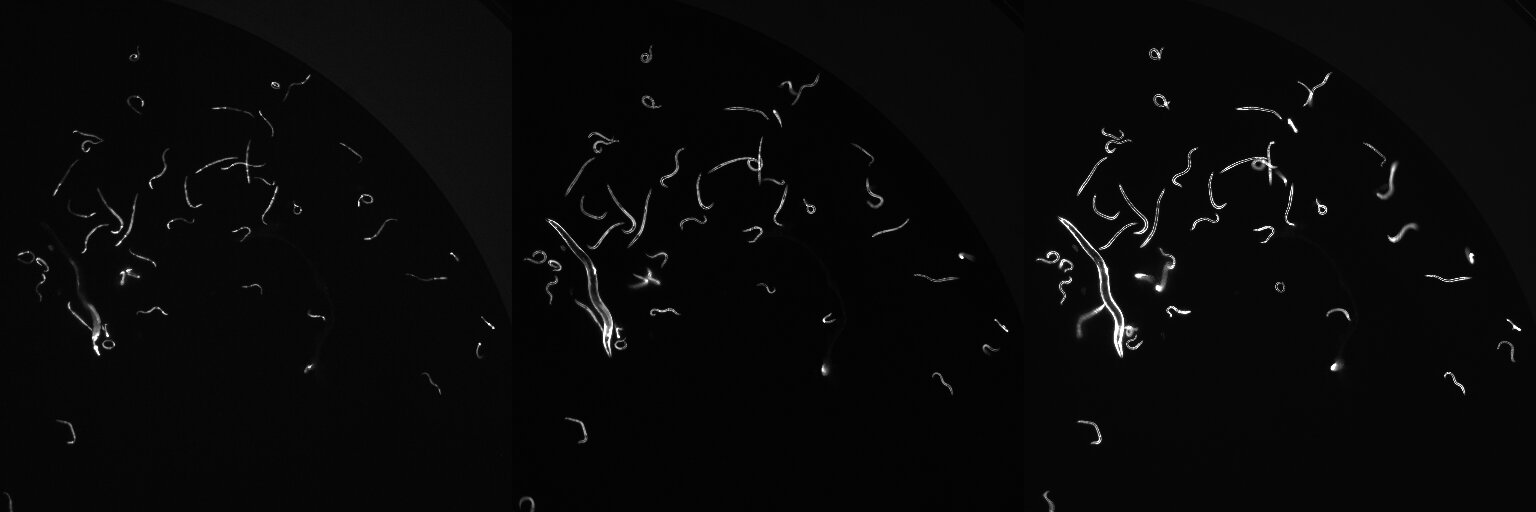

### Supplemental Figure 3

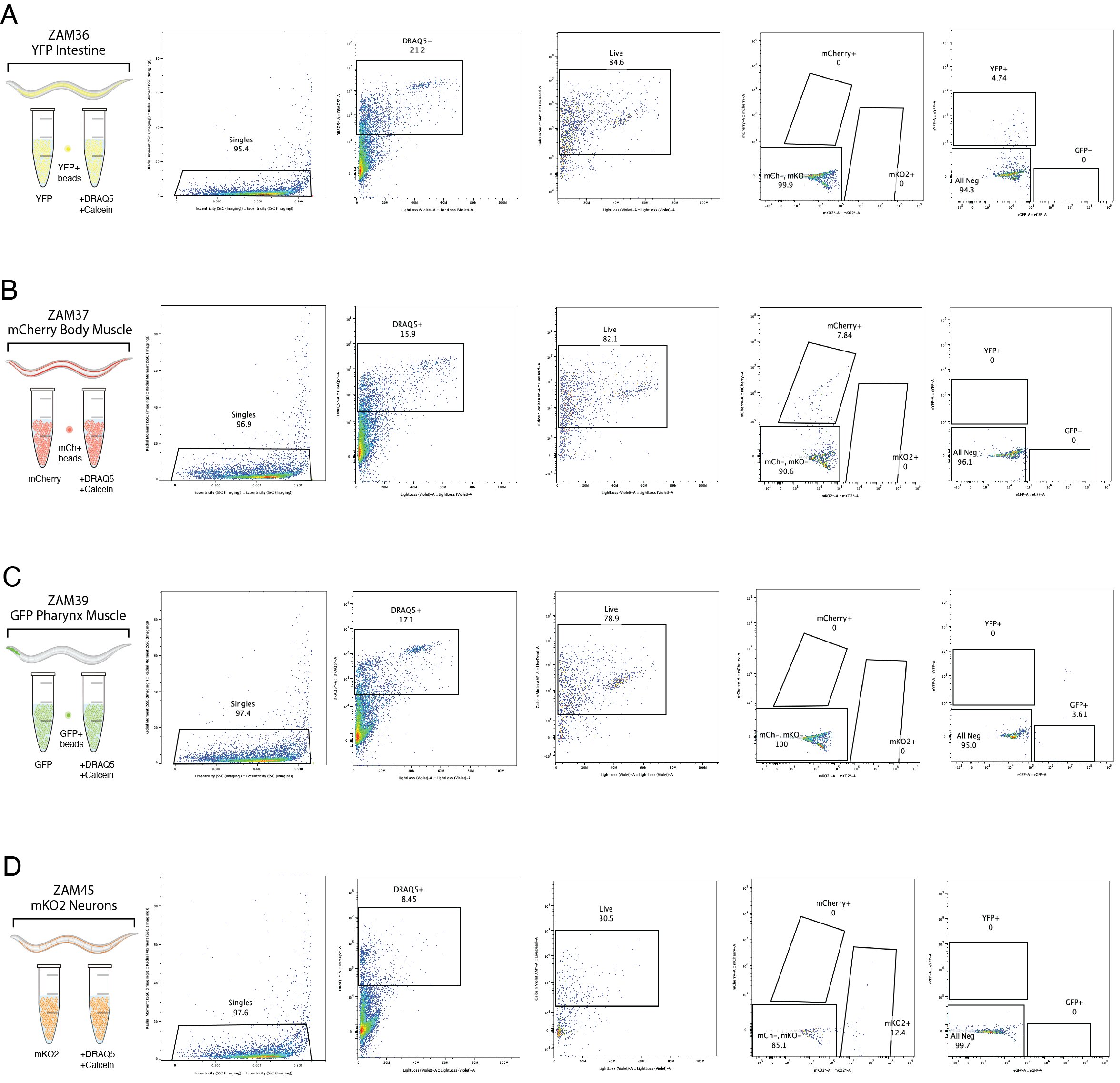
